# LNX1 Modulates Notch1 Signaling to Promote Expansion of the Glioma Stem Cell Population During Temozolomide Therapy in Glioblastoma

**DOI:** 10.1101/2020.09.10.287904

**Authors:** Shivani Baisiwala, Robert H Hall, Miranda R Saathoff, Jack M Shireman, Cheol Park, Louisa Warnke, Clare Hardiman, Jenny Y Wang, Chirag Goel, Shreya Budhiraja, Kathleen McCortney, Craig M. Horbinski, Atique U. Ahmed

## Abstract

Glioblastoma (GBM) is the most common primary brain malignancy in adults, with a 100% recurrence rate and a 21-month median survival. Our lab and others have shown that GBM contains a subpopulation of glioma stem cells (GSCs) that expand with chemotherapy, and that may contribute to therapeutic resistance and recurrence in GBM. To investigate the mechanism behind this expansion, we applied gene set expression analysis (GSEA) to patient-derived xenograft (PDX) cells in response to temozolomide (TMZ), the most commonly used chemotherapy against GBM. Results showed significant enrichment of cancer stem cell and cell cycle pathways (FDR<0.25). The ligand of numb protein 1 (LNX1), a known regulator of Notch signaling by targeting negative regulator Numb, is strongly upregulated after TMZ therapy (p<0.0001) and is negatively correlated with survival of GBM patients. LNX1 is also upregulated after TMZ therapy in multiple PDX lines with concomitant downregulations in Numb and upregulations in intracellular Notch1 (NICD). Overexpression of LNX1 results in Notch1 signaling activation and increased CSC populations. In contrast, knocking down LNX1 reverses these changes, causing a significant downregulation of NICD, eliminating induction of functional stemness after TMZ therapy, and resulting in more prolonged median survival in a mouse model. Our data indicate that removing LNX1 activity results in a less aggressive and more chemo-sensitive tumor. Based on this, we propose that during anti-GBM chemotherapy, LNX1-regulated Notch1 signaling promotes stemness and contributes to therapeutic resistance.

## INTRODUCTION

Glioblastoma (GBM) is an extremely aggressive primary brain malignancy in adults. It has a close to a 100% fatality rate, largely due to an almost inevitable regrowth of the tumor after treatment [1, 2]. The current standard of care is a combination of surgical resection, tumor treating fields, temozolomide (TMZ)-based chemotherapy, and radiation treatment. However, even with that standard of care, patients have a median survival of approximately 21 months with significant decline in their quality of life throughout the treatment regimen [3-5]. The incredibly low survival time is driven by the fact that the recurrent tumor is resistant to our current therapeutic strategies, necessitating the investigating the mechanism of therapeutic resistance.

It is now well-established that GBM contains a subpopulation of cells known as glioma stem cells (GSCs) that contribute to the therapeutic resistance displayed by the recurrent tumor [6-10]. We and others have demonstrated that, during therapy, expansion of the treatment-resistant GSC pool appears to play a crucial role in therapeutic resistance [11, 12]. In order to prevent this expansion, it is necessary to understand exactly how GSC populations are enriched during and after treatment. Previous studies have suggested that GSCs expand through both symmetric and asymmetric cell division [13, 14]. Symmetric cell division is when one GSC gives rise to two more GSCs, whereas asymmetric cell division is when a GSC gives rise to one GSC and one differentiated cell. However, it is not clear if cell division patterns play a role in post-therapy stemness in GBM. Here, we examine if anti-GBM chemotherapy TMZ influences polarized cell division to enhance the post-therapy cellular plasticity in GBM [14].

Notch1 has consistently been shown to play a role in cell division and stemness, and thus in therapeutic resistance, both in GBM and in other cancers [15-19]. Unfortunately, clinical trials directly targeting Notch1 have consistently failed due to unacceptable side effects from Notch1 inhibitors [20]. Here, we examine LNX1 as a regulator of Notch1 signaling to ultimately control expansion of the stem cell compartment after TMZ therapy. This allows us a novel pathway through which to target oncogenic activity of Notch1 in the GBM – a pathway that is much more targeted to tumor cells and therefore may be much less likely to be toxic to patients.

## MATERIALS & METHODS

### Gene Set Enrichment Analysis

RNA was extracted from samples using the RNEasy kit (Qiagen, Hilden, Germany), per manufacturer’s instructions. For each analysis, 1ug of RNA was utilized with Human HT12 (48,000 probes, RefSeq + EST), again per manufacturer’s directions (Illumina, San Diego, CA, USA). All microarrays were performed in triplicate to ensure appropriate replicates for the experiment.

This data was analyzed for gene expression pre- and post-TMZ, looking at genes involved in cell division and stemness pathways. This work was performed by a trained biostatistician using an established gene set enrichment analysis algorithm [21]. Briefly, this algorithm identifies biological pathways as a whole and assesses their enrichment over different conditions as well as the significance of that enrichment (FDR value). It also generates fold change for expression for specific genes based on the microarray data.

### RNA-Seq Analysis

Cells were treated for four days with 50 µM TMZ or equimolar DMSO in triplicate. Cell pellets were then harvested and are sent to Novogene’s sequencing lab (UC Davis, CA, USA). Novogene performed extractions and sequencing and returned results of sequencing.

This data was analyzed for gene expression pre- and post-TMZ, looking at genes involved in polarized cell division, as identified in *Drosophila* studies [14]. This work was performed by applying the DESeq2 algorithm as previously described across our genes of interest [22]. Briefly, this algorithm estimates the dispersion of gene counts as compared to an overall mean and thus generates a fold change and a p-value for each gene of interest.

### Cell Lines and Culture

U251, a human glioma cell line, was obtained from the American Type Culture Collection (Manassas, VA, USA). Cells were cultured in Dulbecco’s Modified Eagle’s Medium (DMEM; HyClone, Thermo Fisher Scientific, San Jose, CA, USA), with 10% fetal bovine serum (FBS; Atlanta Biologicals, Lawrenceville, GA, USA) and 1% penicillin-streptomycin antibiotic mixture (Cellgro, Herndon, VA, USA; Mediatech, Herndon, VA, USA).

Our patient-derived xenograft (PDX) glioma cells (GBM43, GBM5, and GBM6) were obtained from Dr. C. David James at Northwestern University and maintained according to published protocols [23]. They were cultured in DMEM, supplemented with 1% FBS and 1% penicillin-streptomycin. Cells were used for a maximum of 4 passages before being replenished from frozen stock. Frozen cells were maintained in pure FBS supplemented with 10% dimethyl sulfoxide (DMSO) in liquid nitrogen at −180°C.

### Immunofluorescence Staining

For *in vitro* experiments, cells were cultured as described above. In preparation for staining, they were re-plated in Lab-Tek II 8-well chambers (Thermo Fisher Scientific, Waltham, MA, USA) and then treated with the appropriate drug (DMSO or TMZ 50 µM) 24 hours after being plated. After treatment duration was complete, cells were washed once with PBS and then fixed for 10 minutes at room temperature with cold 4% PFA (Thermo Fisher Scientific, Waltham, MA, USA). They were washed after fixation and then blocked in 10% BSA + Triton-X (Thermo Fisher Scientific, Waltham, MA, USA) for 2 hours. Cells were then maintained in the appropriate primary antibodies – LNX1 1:200 (Invitrogen, Carlsbad, CA) or Numb 1:200 (Thermo Fisher Scientific, Waltham, MA, USA) – diluted in 1% BSA overnight at 4°C. The next day, cells were washed 3 times for 8 minutes each and then placed in the secondary antibodies conjugated to either Cy3 or FITC (Thermo Fisher Scientific, Waltham, MA, USA) at a dilution of 1:1000 in 1% BSA for 2 hours. Following secondary incubation, cells were washed 3 times for 10 minutes and then mounted with DAPi containing mounting media (Invitrogen, Carlsbad, CA, USA).

For *in vivo* experiments, mice were perfused with ice-cold PBS (Gibco, Waltham, Massachusetts, USA) following euthanasia. They were then frozen in cryoprotectant on ethanol and dry ice and were subsequently stored at −80°C until required for experimentation. For staining, samples were sectioned at a thickness of 8 uM and then stained per standard immunohistochemistry protocols [24]. In brief, sections were thawed for 30 minutes at 37°C. They were washed with PBS 3 times for 5 minutes each following thawing to remove additional cryoprotectant. The cells were then fixed in cold 4% PFA (Thermo Fisher Scientific, Waltham, MA, USA) at room temperature for 15 minutes and were once again washed with PBS 3 times for 5 minutes each. Sections were then incubated in 10% BSA + Triton-X for 2 hours to block and then incubated in the appropriate primary antibodies as above overnight at 4°C. Following this, sections were washed in PBS 3 times for 10 minutes each and then incubated in secondary antibodies as above at 1:1000 for 2 hours at room temperature. Finally, samples are thoroughly washed (3-4 times for 10 minutes each) and mounted with DAPI-containing mounting media.

All slides were imaged using a Leica confocal and images were analyzed using ImageJ software [25]. Where relevant, counts of cell division were performed by calculating relative intensity values in ImageJ. In addition, where relevant, co-localization correlation coefficients (i.e. Pearson’s correlation coefficient) were calculated by using in-built Coloc2 package in ImageJ [26].

### Immunohistochemistry of Human Samples

Human glioma samples (primary and recurrent) are obtained from Northwestern University’s Nervous System Tumor Bank. All patients gave consent per the defined Institutional Review Board (IRB) policies prior to obtaining samples. Staining was performed per standard immunohistochemistry protocols, previously established in that lab [24]. Briefly, samples were formalin-fixed and paraffin-embedded (FFPE). They were then sectioned at a thickness of 4uM, after being heated at 60°C for at least 1 hour. Antigen retrieval was performed with a BioCare Medical Decloaking Chamber using high (LC3) or low pH antigen retrieval buffer from Dako (Agilent, Santa Clara, CA, USA). Primary antibodies were incubated for 1 hour at room temperature followed with horseradish peroxidase (HRP)-tagged secondary antibodies as appropriate. Slides were scored for LNX1 expression on a scale of 1 (lowest) to 3 (highest) by a board-certified neuropathologist (CMH), and scores were plotted alongside survival data.

### Quantitative PCR

Cells were harvested and RNA extraction was performed using Qiagen RNA extraction kits (Qiagen Hilden, Germany), per manufacturer’s protocol. cDNA was generated from RNA samples using iScript kits (BioRad, Hercules, CA, USA), per manufacturer’s protocol. Once generated, cDNA was diluted 1:10 to use for downstream qPCR. PCR reactions were set up with a standard amount of cDNA, SyberGreen, forward and reverse primers, and ddH2O. All primers were generated from Primer-BLAST using the native settings. Reactions were all performed in biological triplicate and technical duplicate.

### Western Blot

Cells were harvested and protein was extracted using mPER lysis buffer (Thermo Fisher Scientific, Waltham, MA, USA) with protease and phosphatase inhibitors (Cell Signaling Technologies, Danvers, MA, USA). Cells were then sonicated 3 times for 30 seconds each with 10 second intervals in between. Next, they were incubated on ice for 10 minutes and then centrifuged for 10 minutes at 13,000 rpm. The clear supernatant was recovered and used for protein assays. Samples were equalized using the Pierce BSA Protein Assay Kit (Thermo Fisher Scientific, Waltham, MA, USA), per manufacturer’s instructions. Prepared samples were incubated at 95°C for 10 minutes and were then cooled to room temperature and loaded into a pre-poured 10% SDS-PAGE gel. Gels were run for 30 minutes at 40V and then 95V until samples had run all the way through. Gels were then transferred to nitrocellulose membranes for 1 hour at 14V, blocked for an hour in 5% milk (Thermo Fisher Scientific, Waltham, MA, USA), and incubated overnight in primary antibody diluted in 5% BSA (Thermo Fisher Scientific, Waltham, MA, USA). Primaries used include LNX1 1:1000 (LSBio, Seattle, WA, USA) and Numb, Notch1, Hes1, p21, MAML, beta-Actin all used at a dilution of 1:1000 (Cell Signaling Technologies, Danvers, MA, USA). The next morning, membranes were washed 3 times for 10 minutes in TBS-T buffer and then incubated in secondary antibodies made against mouse and rabbit as appropriate in milk at a dilution of 1:4000 (Cell Signaling Technologies, Danvers, MA, USA). Membranes were washed again 3 times for 20 minutes and then developed using the BioRad imaging system (BioRad, Hercules, CA, USA).

If immunoprecipitation was required prior to western blotting, cells were thoroughly washed and then protein was isolated using mPER and protease/phosphatase inhibitors as described above. However, prior to equalization and further blotting, an immunoprecipitation was performed using the antibody of interest and the Protein A/G Plus Ultralink Resin Kit (Thermo Fisher Scientific, Waltham, MA, USA), per manufacturer’s instructions. Briefly, the protein extract and relevant antibody were incubated together overnight with gentle rotation at 4°C. The next day, a coupling with UltraLink resin was performed for 2 hours at room temperature, also with gentle rotation. Following that, IP buffer was added, and samples were centrifuged for 3 minutes at 2500g, after which the supernatant was discarded. This step was repeated 3 times. Finally, the binding protein was eluted with 50uL of elution buffer and western blot samples were then prepared as above.

### Neurosphere Assays & Extreme Limiting Dilution Analysis (ELDA)

Cell lines were cultured as described. They were then harvested, washed with PBS, and plated in serial dilutions, specifically 200, 150, 100, 50, 25, 12, 6, and 3 cells per well. Each dilution was performed in 12 replicates. Cells were maintained in neurobasal media (Gibco, Waltham, MA, USA) supplemented with B27 (no Vitamin A; Invitrogen, Carlsbad, CA, USA), basic fibroblast growth factor (bFGF; 10ng/ml; Invitrogen, Carlsbad, CA, USA), epidermal growth factor (EGF; 10ng/mL; Invitrogen, Carlsbad, CA, USA), and N2 (Invitrogen, Carlsbad, CA, USA). Cells were treated either with 50 uM TMZ or equimolar DMSO. A blinded experimenter examined the wells after 7 days. The number of formed neurospheres with a diameter greater than 20 cells was counted. Counts were analyzed using the Walter + Eliza Hall Institute of Medical Research platform (http://bioinf.wehi.edu.au/software/elda/). This platform allows for determination of stem cell frequency and quantification of significant differences between groups. In addition, the absolute number of spheres was plotted visually, and images were taken of the wells using a Leica confocal microscope.

### Notch Reporter Transfection

Cells were plated at approximately 60% confluency, 24 hours prior to transfection. HP DNA transfection reagent (Sigma Aldrich, St Louis, MO, USA) was used to perform transfections, per manufacturer’s protocols. Briefly, transfection reagent and Notch reporter plasmids (#44211, 47684, 47683; Addgene, Cambridge, MA, USA) were combined in serum free OptiMEM media (Gibco, Waltham, MA, USA). Specifically, 2 ug of plasmid was used per well for a 6-well plate and transfection reagent was added at a ratio of 1:3. The mix was incubated for 30 minutes with a thorough vortex every 10 minutes. Then, it was distributed dropwise to plated cells. Cells were maintained for 48-72 hours and were then harvested for downstream analysis by flow cytometry, as described below.

### Flow Cytometry

Cells were cultured as described above. At relevant time points following transfection or transduction as needed, cells were collected, and fresh surface staining was performed. Cells were first washed with sterile PBS (Gibco, Waltham, Massachusetts, USA). Next, they were detached from the plate using 0.05% trypsin / 0.53mM EDTA (Corning, Corning, New York, USA). Trypsin was neutralized using an equal amount of culture media, and cells were collected in appropriately sized tubes. Cells were then incubated with conjugated antibodies against CD133-APC (Miltenyi Biotc, Bergisch Gladbach, Germany) for 30 minutes at room temperature. They were washed thoroughly with PBS (3 washes for 5 minutes each) and were analyzed using the flow cytometer. Further analysis and quantification of results was performed using the FlowJo software.

### Viability Assays

Viability assays were conducted using the MTT assay. In brief, cells were plated at a density of 3,000 cells per well in a 96-well plate with 8 replicates plated for each condition. After 24 hours, media was changed to media supplemented with the appropriate drug concentration. Three days later, cells were thoroughly washed with phosphate buffered saline (PBS) and incubated with the MTT reagent (Thermo Fisher Scientific, Waltham, MA, USA) diluted 10% in regular culture media. Cells were then maintained for 4 hours at 37°C. After this period, the MTT reagent was removed and samples were thoroughly re-suspended in DMSO. The absorbance of each well was assessed using a standard plate reader and results were tabulated per standard protocols [27].

### Generation of Viral Particles

Low passage 293T cells (ATCC, Manassas, VA, USA) were used to generate lentiviral particles. Briefly, cells were plated at 90% confluency in preparation. After 6 hours, they were transfected with a mix of HP DNA Transfection Reagent (Sigma Aldrich, St Louis, MO, USA) diluted in OptiMEM media (Gibco, Waltham, MA, USA) as well as appropriate packaging plasmids and CRISPR-Cas9 plasmids, per manufacturer’s instructions. Inducible Cas9 was obtained from Addgene (Cambridge, MA, USA) and guide RNA plasmids were obtained from Genecopoeia (Rockville, MD, USA). The transfected 293T cells were maintained in culture for 48-72 hours, after which the virus-containing supernatant was harvested. The supernatant was centrifuged at 1200rpm for 5 minutes to remove cell debris. It was additionally filtered with a 45-micron filter to sterilize. It was then ultra-centrifuged at 288,000g for 3 hours to pellet virus particles. Particles were resuspended in 200ul of PBS and frozen at −80°C until use.

### Transduction of Cell Lines with Lentiviral Particles

U251 and GBM43 lines were obtained and maintained in culture as detailed above. For infection, cells were harvested and resuspended in a small volume of media (∼50 ul). Appropriate lentivirus amounts were added (∼10-20 MOI per sample). Polybrene (4ug/mL) was added to the virus-cell mixture. The tubes were incubated for 30 minutes at room temperature and were then plated in appropriately sized flasks. Cells were maintained in culture with regular media changes for 48-72 hours. Efficiency of the resulting modifications was assessed by western blotting, as previously described.

### Animal Studies

Mice used in this study are athymic nude mice (nu/nu; Charles River, Skokie, IL, USA). They were housed in accordance with all Institutional Animal Care and Use Committee (IACUC) requirements and in compliance with all applicable federal and state statutes. Animals were housed in shoebox cages with food and water available and with a strict 12-hour light and dark cycle.

Our lab has a previously established glioblastoma mouse model, where intracranial implantation of glioblastoma cells is performed [28]. Briefly, animals received an injection of buprenex and metacam by intraperitoneal (IP) injection. Next, they received a second injection of ketamine/xylazine anesthesia (Henry Schein; New York, NY, USA). Complete sedation of the mice was confirmed by pinching the foot. To protect the mice, artificial tears were then applied to each eye and the scalp was sterilized with ethanol and betadine. A small incision was made using a scalpel, exposing the skull. A drill was used to make a ∼1mm burr hole above the right frontal lobe. A stereotactic rig and a Hamilton syringe loaded with cells were used to implant 5 uL of cell solution 3 mm from the dura. Injections occurred over a period of one minute. The needle was then raised slightly and left for an additional one minute to ensure release of the cell suspension. Finally, the syringe was carefully removed, and the scalp was closed with sutures (Ethicon; Cincinnati, OH, USA). Head position was maintained throughout the closing process. Animals were maintained on heat pads until awake and reactive following surgery.

Any drug treatments necessary were started one week following the implantation. Animals received IP injections of either TMZ (2.5 mg/kg) or equimolar DMSO for 5 consecutive days, once per day. Animals were monitored daily for any signs of tumor progression (i.e. weight reduction, reduced body temperature, hunching, etc.). Animals were euthanized when it was determined that they would likely not survive to the next morning by two independent researchers. These sacrifices were performed according to Northwestern University and IACUC guidelines.

### Statistics

Statistical analyses were performed with GraphPad Prism v8 (GraphPad Software; San Diego, CA, USA). Data are presented as mean for continuous variables and number or percentage for categorical variables. Differences between two groups were assessed using Student’s t-test. Differences between multiple groups were assessed using analysis of variance (ANOVA) with Turkey’s post-hoc correction. Survival curves were graphed with the Kaplan-Meier method and compared by log-rank test. All tests are two-sided and a p-value of under 0.05 is considered significant for the purposes of our study.

## RESULTS

### LNX1 Expression and Symmetric Cell Division Are Enriched After TMZ-Based Chemotherapy

To investigate the mechanism of chemoresistance in GBM, we performed a gene expression analysis during TMZ therapy in a treatment-naïve patient-derived xenograft (PDX) model. Cells were treated with the physiologically relevant dose of TMZ (50 µM) or with equimolar vehicle control DMSO [12]. After 4 days of treatment, cells were harvested and analyzed for gene expression. Enrichment analysis (GSEA) revealed significant enrichment for stemness [28], cell cycle (*FDR = 0*.*0001*), and hypoxic response (*FDR = 0*.*015*) gene networks after therapy with TMZ indicating that cells are altering their division patterns and upregulating their stemness phenotype upon treatment with TMZ (**Fig 1A**).

**Figure 1:**
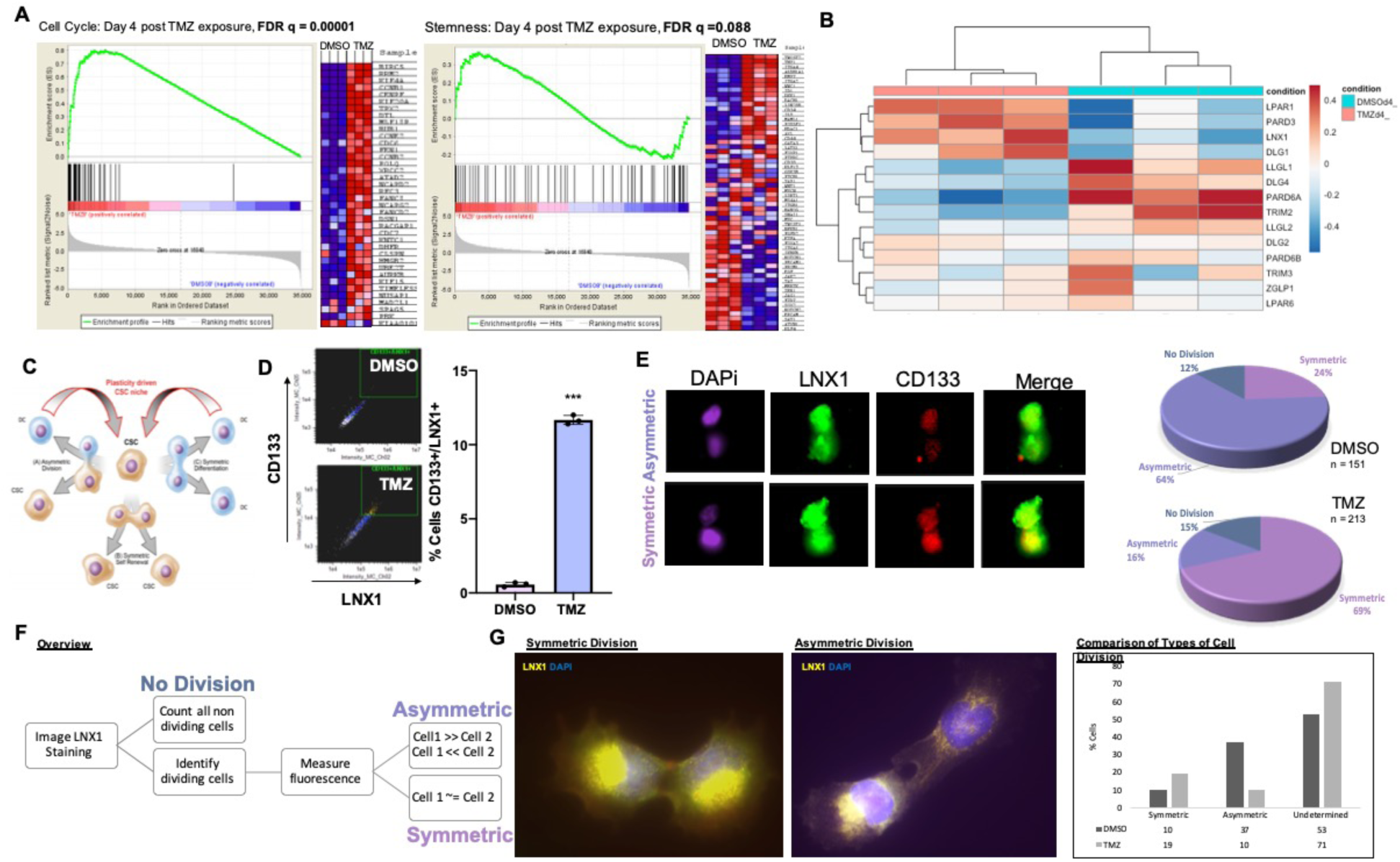
Polarized LNX1 Expression during TMZ therapy. (A) GSEA analysis comparing GBM43 cells treated for 4 days with DMSO or 50 µM TMZ showed significant enrichment of cell division and stemness pathways. (B) RNA-Seq analysis of a panel of genes involved in polarized cell division in GBM43 cells treated with DMSO vs 50 µM TMZ showed significant enrichment of LNX1 expression after TMZ therapy. (C) Stem cell division in cells can produce either 2 differentiated cells, 2 stem cells, or 1 of each. (D) Analysis of GBM6 cells by ImageStreamX technology revealed that expression of LNX1/CD133 significantly increased after TMZ therapy. (E) Image analysis showed that LNX1 and CD133 reliably colocalize and that TMZ therapy significantly increases the percentage of symmetric stem cell divisions as compared to DMSO. (F&G) U251 cells were analyzed manually in an experiment comparing dividing cells treated with DMSO vs 50 µM TMZ. Similar results to prior were observed, where TMZ significantly increased the proportion of symmetrically dividing stem cells as compared to control. Data were analyzed using ImageStreamX software or Prism 8, as appropriate. Student’s t-test was used to calculate significance between 2 groups. ANOVA was used for greater than 2 groups. *p < 0.05; **p < 0.01; ***p < 0.001; ns, not significant

Our lab, along with others, has previously demonstrated that stresses generated by chemo- and radiotherapy can promote expansion of the GSC population by differentiated GBM cells dedifferentiating and acquiring a stem-like phenotype [12, 29, 30]. Previous reports also demonstrate that GSCs are capable of maintaining stemness in GBM through symmetric cell division [8]. Further, in low-grade oligodendrogliomas, defects in asymmetric cell division are associated with the neoplastic transformation [14]. These reports raise the possibility that cell polarity may contribute to the post-therapy expansion of the GSC subpopulation and promote therapeutic resistance. To examine this, we first analyzed gene expression patterns for a panel of known genes controlling asymmetric vs. symmetric cell division [13, 31-33]. The RNA-Seq experiment was first analyzed to assess sample similarity and screen quality (**Fig S1**). The expression of LPAR1, PARD3, LNX1 and DLG1 were approximately three- to four-fold elevated as compared to other genes, after treatment with TMZ (**Fig 1B**).

LNX1 is an E3 ubiquitin ligase that is thought to promote symmetric cell division of stem cells by acting through Numb to regulate Notch signaling [34]. To further investigate this in glioma stem cells (GSCs) specifically during therapy, we used Image-Stream-X analysis, which allows for simultaneous cell imaging and flow cytometry. These images could then be analyzed for cell division patterns and categorized into symmetric self-renewal, symmetric differentiation, or asymmetric division (**Fig 1C**). Cells were treated with DMSO or TMZ for 4 days before being analyzed by ImageStreamX technology. Flow cytometry analysis of the cells showed that LNX1 and CD133 expression were definitively colocalized after TMZ therapy (*p < 0*.*001*, **Fig 1D**). Furthermore, visual analysis of these images revealed that LNX1 had a propensity to be segregated with CD133 in dividing cells. As CD133 is a phenotypic marker of GSCs, these together provide evidence that segregation of LNX1 is associated with possible maintenance of stemness phenotype during TMZ therapy [35]. Quantitative analysis of LNX1/CD133 expression showed that cells exposed to TMZ significantly upregulated the proportion of symmetric expression of LNX1/CD133 expression, thus suggesting the possibility that expansion of the GSC compartment is through symmetric division in response to TMZ exposure (**Fig 1E**).

In order to further confirm our Image-Stream-X result using another experiment, we performed immunofluorescence in PDX GBM cells for LNX1 after treating cells with TMZ for 4 days. The cells were imaged, and the fluorescence intensity of LNX1 was calculated for all sets of dividing cells, as identified by phalloidin staining. Dividing cells that had a fluorescence ratio of greater than 1.5 were considered to be dividing asymmetrically while the rest were classified as dividing symmetrically (**Fig 1E**). This ratio was determined based on other similar studies [35]. Those cells that were not dividing or could not be measured accurately were classified as indeterminate. This analysis also showed that the proportion of symmetric cell division is increased after therapy with TMZ, reflecting that TMZ therapy both increases LNX1 expression and promotes a concurrent increase in the percentage of symmetric segregation of LNX1 expression, thereby possibly contributing to the expansion of the GSC compartment during TMZ therapy (**Fig 1F**).

### LNX1 Expression Is Elevated in Post-Therapy Recurrent GBM And Negatively Correlated with Survival with Patients With GBM

Our next step was to examine the clinical significance of LNX1 in gliomagenesis by examining various GBM patient datasets as well as GBM tissue. Analysis of the Cancer Genome Atlas (TCGA) data through the GlioVis portal showed that LNX1 mRNA expression increases progressively with GBM recurrence (**Fig 2A**). In addition, it was negatively correlated with patient survival in two different datasets (**Fig 2B**) [36]. We next validated these results by further examining existing publicly available GBM patient datasets. Analysis of TCGA data through the cBioPortal interface showed that LNX1 was amplified in 10% of patients, independent of their IDH-1 status (TCGA Cell 2013, 543 samples; **Fig 2C**) [37, 38]. Finally, to investigate the LNX1 protein expression in the GBM tissue, we performed immunohistochemistry for LNX1 on de-identified patient samples from consenting patients at Northwestern Medicine. The sections were then scored for LNX1 presence within the tumor body. A score of 1 corresponded to low expression, while a score of 3 corresponded to high expression (**Fig S2**). These scores were then plotted alongside patient survival (**Fig 2D**). The analysis showed that an increase in LNX1 expression scores corresponded to a decrease in patient survival (*p = 0*.*04*).

**Figure 2:**
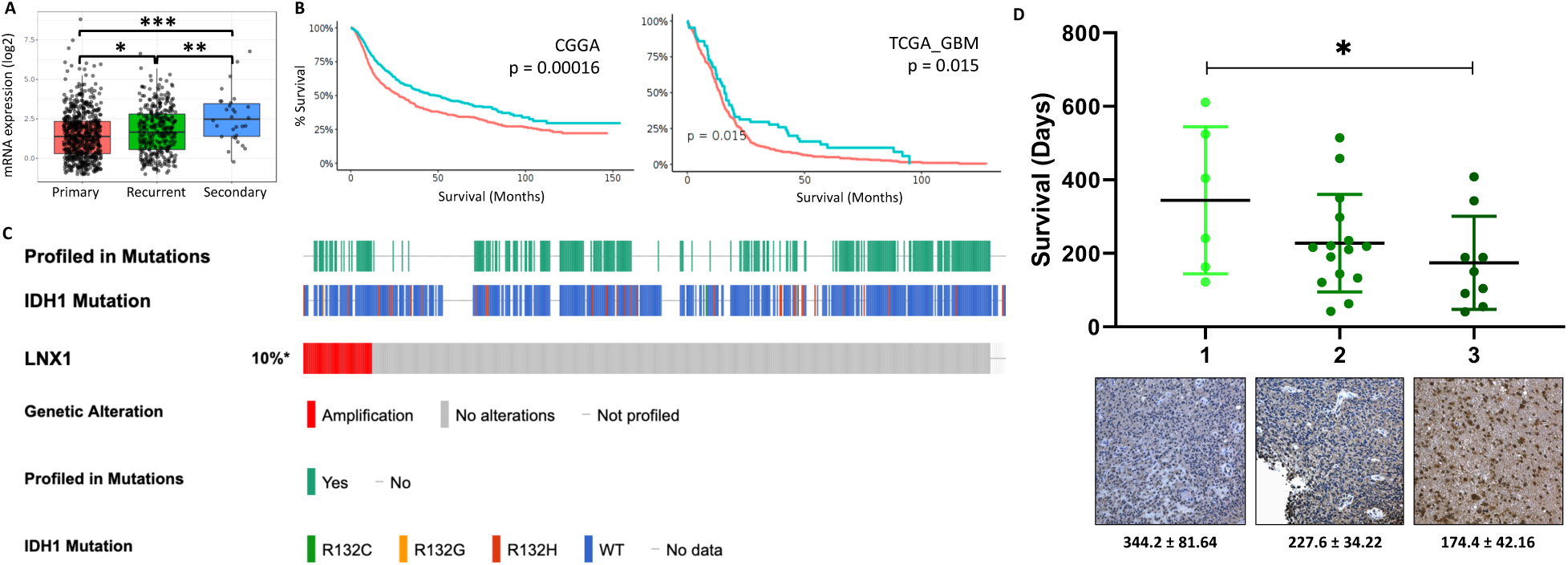
LNX1 is Elevated in GBM Patients and Correlates with Patient Outcomes. (A) LNX1 expression was approximated in the CGGA dataset and was shown to elevate significantly with each recurrence (B) In two datasets, using optimal cutoff analyses, elevated LNX1 expression was shown to be correlated with significantly worsened survivals for patients. (C) LNX1 expression was tabulated across over 500 patient samples, based on TCGA data, and showed amplification in 10% of samples, with no correlation to the IDH status of the patient. (D) LNX1 expression was scored in patient samples by a certified neuropathologist and scores were plotted against survival, with a significant difference in survival between high and low LNX1 expression. Significance was calculated in Prism 8 using Student’s t-test for 2 samples or ANOVA for greater than 2 samples. Tukey’s Honest Significance Difference was applied to determine significance for TCGA expression data and log-rank testing was used to compare survival curves. *p < 0.05; **p < 0.01; ***p < 0.001; ns, not significant.

### LNX1 Influences Notch1 Expression Through an LNX1-Numb-Notch1 Axis During TMZ Therapy

Multiple studies have shown that LNX1, which is an E3 ubiquitin ligase, acts on Numb in the cell by ubiquitinating Numb and tagging it for degradation [37, 38]. Furthermore, many other papers have focused on the importance of Numb in modulating Notch1 activity and thus further modulating downstream proteins, especially for regulation of cell proliferation and expansion [39]. As Notch has been established as a critical signaling hub for glioma progression, we next set to examine the effect of LNX1 elevation after TMZ therapy on this axis [18].

We treated the PDX cell line GBM43 with 50 µM TMZ and equimolar DMSO vehicle control for 4 and 8 days. We then harvested and processed samples to assess levels of mRNA by quantitative PCR (qPCR). Results showed that LNX1 mRNA levels were elevated in both timepoints post-therapy. We investigated the LNX1 transcript distribution in the different cellular compartments within GBM tissues using the IvyGap dataset, which relies on *in-situ* hybridization to identify expression patterns. We noted a similar pattern here, with LNX1 upregulations associated with Numb downregulations and NICD upregulations (**Fig S4**).

Furthermore, we were able to show that the Notch1 downstream target genes (*RBPSUH, p21*, and *Hes1*) also all had elevated mRNA transcripts following therapy (*p < 0*.*01*; **Fig 3A**). We additionally assessed levels of expression at the protein level after therapy. We used GBM6, a classical subtype with unmethylated MGMT; GBM43, proneural subtype with unmethylated MGMT (resistance to TMZ), and GBM5, a mesenchymal subtype with MGMT methylated (TMZ sensitive) to evaluate the LNX1 expression during TMZ therapy. Analysis of all three subtypes of PDX GBM showed that protein levels for LNX1 increased with a corresponding decrease in Numb levels (**Fig 3C)**. This resulted in an increase in the level of intracellular Notch1 (NICD) as well as the downstream targets of Notch1 – RBPSUH, Hes1, and p21 (**Fig 3B**). These data indicated that during TMZ therapy, LNX1 expression is negatively correlated with Numb and positively correlated with Notch1 signaling.

**Figure 3:**
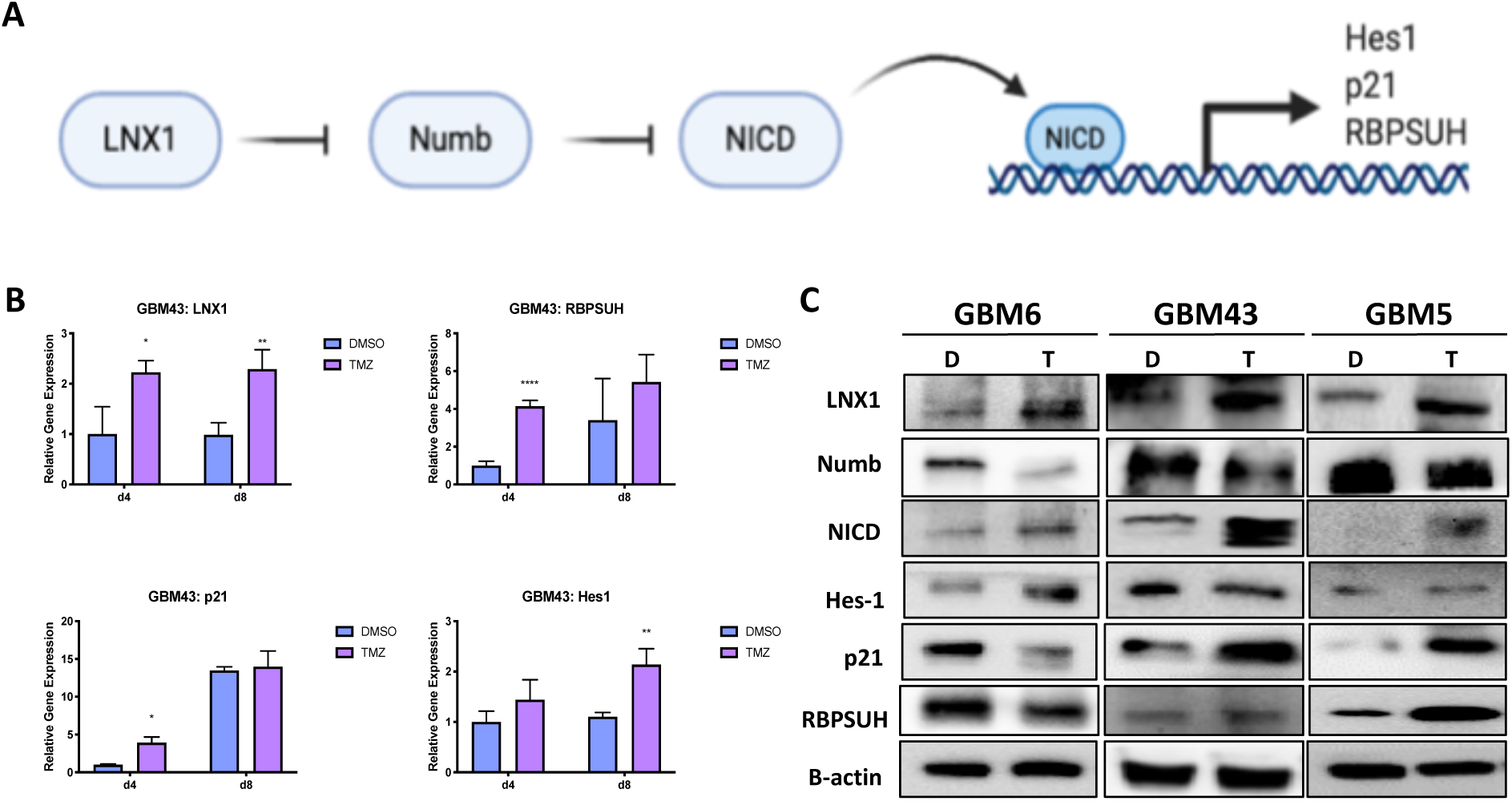
The LNX1-Numb-Notch1 Axis Controls Is Activated After TMZ Therapy in GBM. (A) Examination of mRNA levels 4 and 8 days after therapy with TMZ compared to a DMSO control showed elevations in LNX1, Hes1, p21, and RBPSUH after therapy. (B) Western blotting across multiple GBM PDX cell lines showed that LNX1 was upregulated compared to control after 4 days of TMZ therapy, corresponding with Numb downregulations, and NICD upregulations. Following Notch1 upregulation, Notch1 downstream markers p21, Hes1, and RBPSUH were also elevated. Beta-actin was included as a loading control for all blots performed. Analysis was performed in Prism 8, using ANOVA to compare row-means to determine significance *p < 0.05; **p < 0.01; ***p < 0.001; ns, not significant

### LNX1 interacts with Numb during TMZ therapy

Next, we assessed the interactions between LNX1, Numb, and Notch1 in the cell. In order to do this, TMZ-treated GBM 6 and 43 PDX cell lines were subject to immunoprecipitation western blot (IP-WB) analysis with the antibody of interest to assess levels of interaction between two target proteins. First, we immunoprecipitated samples with an antibody for LNX1. Western blot analysis of these IP samples revealed that there was an increase in binding of Numb and LNX1 in the presence of TMZ, as well as an increase in the ubiquitination of LNX1 in the presence of TMZ. From these results, we inferred that LNX1 is likely transferring ubiquitin to Numb at an increased frequency in the presence of TMZ. A similar analysis with Notch1 pulldown samples showed that Numb-Notch1 interactions were minimally changed following therapy with TMZ (**Fig 4A**). However, ubiquitinated Notch was decreased during TMZ therapy in GBM43, reflecting a subtype-specific differential response. Overall, these IP blots reflect that there is an interaction between LNX1 and Numb and a further interaction between Numb and Notch during therapy.

**Figure 4:**
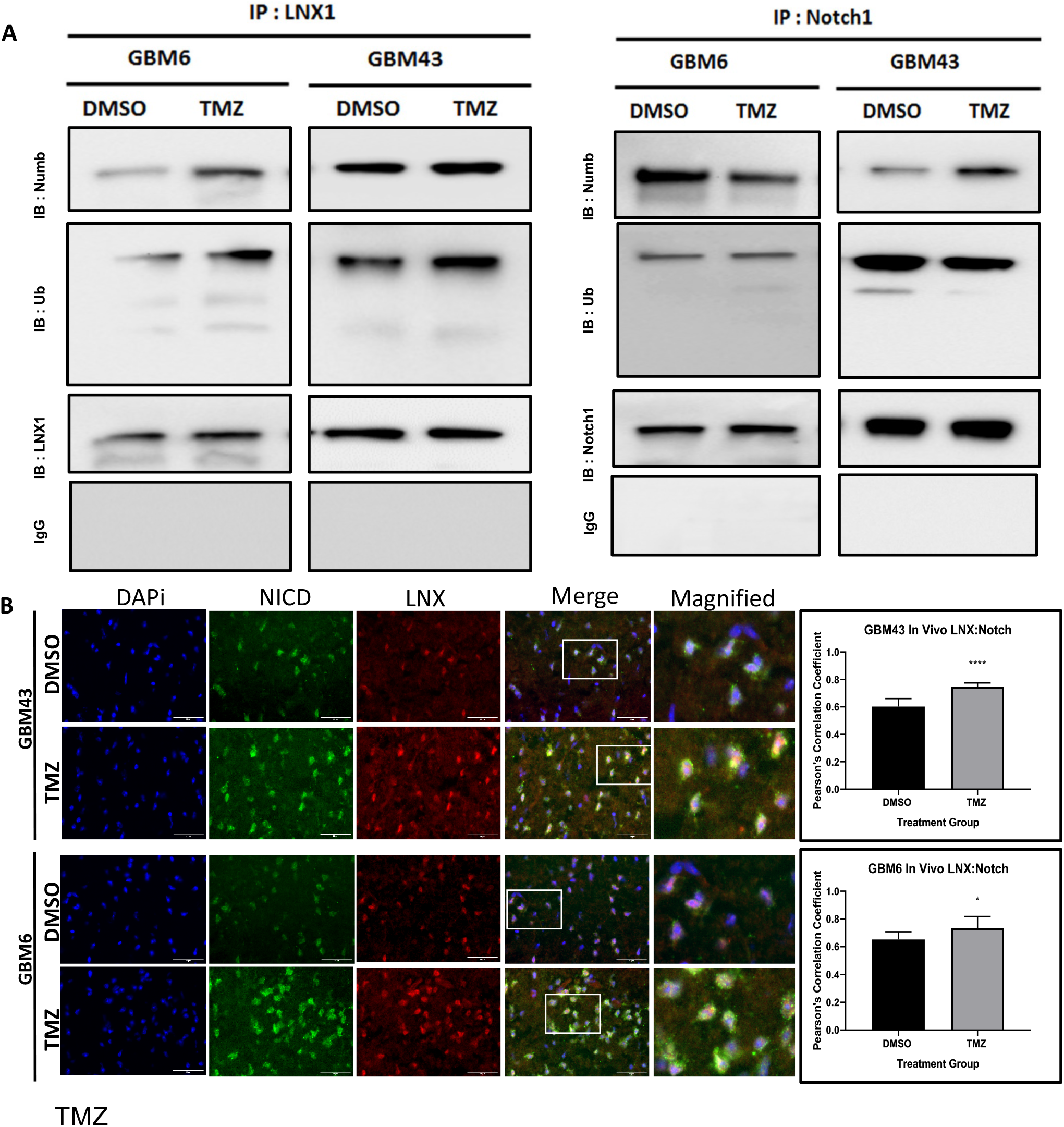
LNX-Numb and Numb-Notch1 interactions Were Upregulated After TMZ Therapy. (A) An IP was performed for LNX1 in GBM6 and GBM43 cells. Samples were then probed for Numb and Ubiquitin. An increase in Numb-LNX interactions and an increase in LNX-Ubiquitin interactions were noted after TMZ therapy, compared to the DMSO control. An additional IP was performed for Notch1 and samples were probed for Numb and ubiquitin levels. A mild increase was noted for Notch1-Numb interactions. IgG controls were performed for all IPs. (B) GBM43 and GBM6 cells were implanted in mice, following our mouse model for intracranial tumors. Tissue samples were isolated from these mice were then stained for LNX1 and Notch1 expression. The tumor area was imaged and colocalization analysis was performed for LNX1 and Notch1 expression. Results showed that LNX1 and NICD colocalization was significantly elevated after TMZ therapy. Image analysis was performed in ImageJ, using the coloc2 package. Statistical analysis was performed in Prism 8, using the Student’s t-test to compare 2 groups and calculate significance. *p < 0.05; **p < 0.01; ***p < 0.001; ns, not significant

To examine this interaction further in vivo, we implanted the GBM43 and GBM6 PDX lines into nu/nu mice, following our previously established tumor mouse model [12]. One week following implantation, mice were treated with 2.5 mg/kg of TMZ for 5 days, delivered once daily. Once the mice reached our clinical endpoint, the animals were sacrificed, and brains were harvested for further experimentation. These brains were sectioned and stained for LNX1 and NICD expression, and images were analyzed for percent colocalization using Pearson’s correlation coefficient. Results showed that therapy with significantly increased LNX1-NICD colocalization in both cell models, further supporting the modulation of Notch1 levels by LNX1 expression (*p <* .*05*; **Fig 4B**).

### Induction of a Stem-Cell Like State Through LNX1 Overexpression Increases Notch1 Activity

Thus far, we have established that LNX1 is a symmetric cell division regulator that is significantly elevated in TMZ. We have further shown that LNX1 acts through interactions with Numb and affects intracellular NICD expression levels. Given that LNX1 regulates Notch signaling, which can further induce expansion of the GSC niche, we hypothesized that elevations in LNX1 during TMZ therapy could be associated with increases in overall stemness of the tumor population [15, 19, 40]. To examine this, we cultured GBM43 and GBM6 PDX lines in neurobasal media to induce a stem-like state and in differentiation media (1% FBS media). We then assessed levels of LNX1, Numb, and Notch1 by western blot. This analysis demonstrated that LNX1 and Notch1 levels are increased when cells are pushed towards a stem-like state and that there is an increased formation of spheres, representative of a stem state, associated with the neurobasal media condition (**Fig 5A, B**). Having observed that LNX1 elevations are indeed associated with the induction of a stem-like state, we wanted to assess the effect of over-expression of LNX1 on the “stemness” of the tumor cell population in the differentiation media. To this end, we subcloned the LNX1 cDNA into an existing lentiviral vector. We then generated lentiviral particles from this plasmid and were able to stably overexpress LNX1 in U251 cells as well as in the GBM43 and GBM6 PDX lines, as validated by western blot analysis. We further noted that Numb expression correspondingly decreased while Notch1 expression increased across all of our cell lines (**Fig 5C**). At the RNA level, gene-level activation of Notch downstream targets RBPSUH, p21, and Hes1 were also observed upon overexpression of LNX1 (*p <* .*05*; **Fig 5D**).

**Figure 5:**
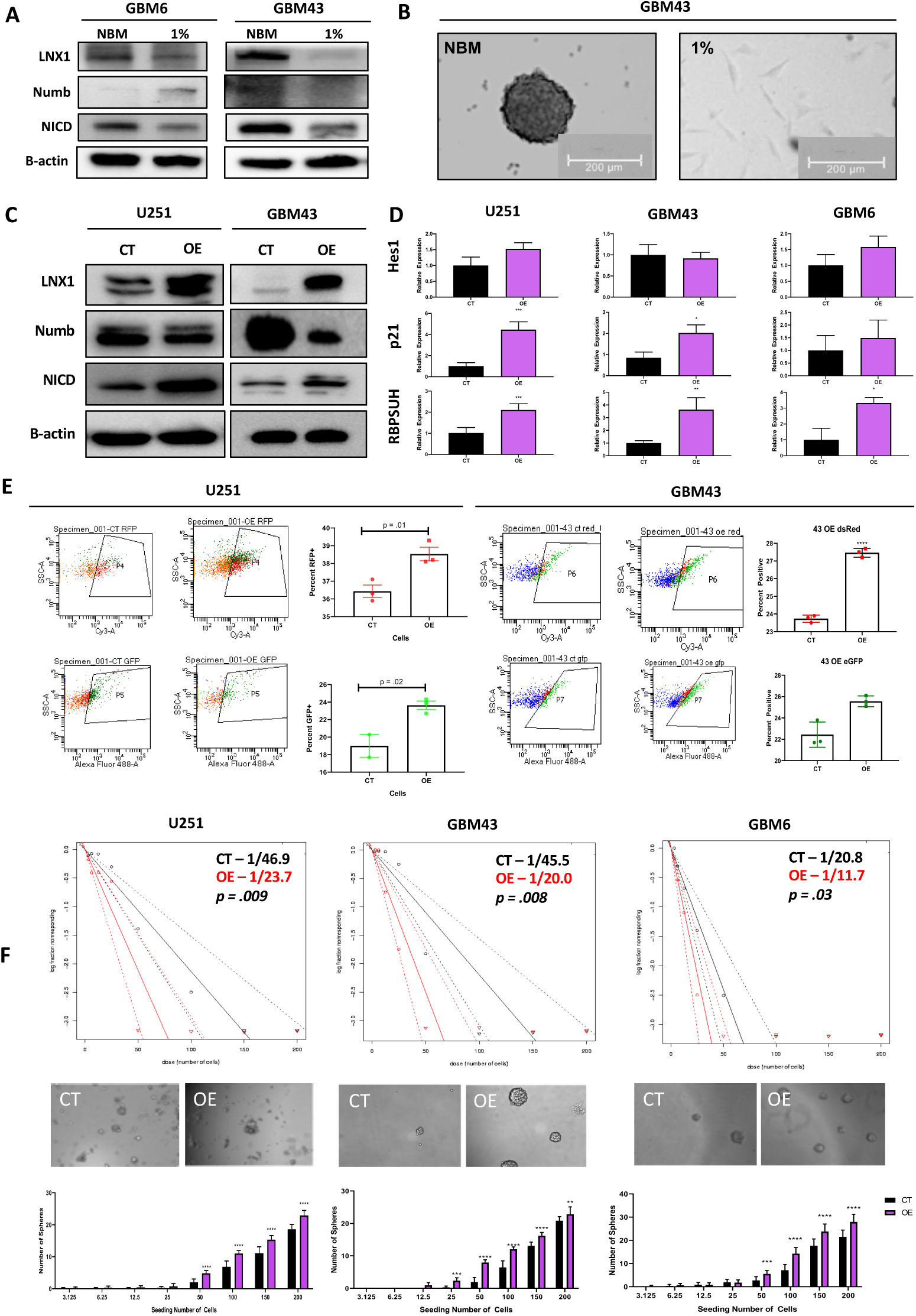
LNX1 Overexpression Results in Notch Activation and Increased Stemness. (A) GBM6 and GBM43 cells are cultured in neurobasal media vs regular 1% media. Neurobasal media showed an increase in LNX1 expression corresponding with a decrease in Numb and an increase in NICD expression. (B) GBM43 cells in neurobasal media demonstrated sphere formation (C) Overexpression of LNX1 in multiple cell lines resulted in decreases in Numb and increases in NICD. (D) NICD increases are matched with an increase in activation of Notch1 downstream markers Hes1, p21, and RBPSUH, measured by quantitative PCR in multiple cell lines. (E) Notch is cleaved to NICD which then trans-locates within the cell and activates transcription, in this case, of our fluorescent reporter proteins. Readout of GFP and RFP reporters separately by FACS shows that LNX overexpression definitively results in activation of Notch1 activity. (F) Limiting dilution neurosphere assays show that LNX1 overexpression results in an increase in functional stemness across multiple cell lines Statistical analysis is performed in Prism 8, using the Student’s t-test to compare 2 groups and calculate significance. Chi-squared testing is used to calculate significance for limiting dilution assays. *p < 0.05; **p < 0.01; ***p < 0.001; ns, not significant

To confirm that Notch1 activation following LNX1 overexpression was functional and could act as a transcription activator, we next used a Notch1 reporter system to read out activity in the LNX overexpressed GBM cells. For this experiment, two separate Notch1 reporters were obtained, each with a different fluorescent tag. It is known that Notch1 is activated by cleavage, and that cleaved Notch1 (NICD) can enter the nucleus and act as a transcription factor. Therefore, these reporter plasmids work by being activated by the cleaved Notch transcription factor to produce either green fluorescent protein (GFP) or red fluorescent protein (RFP). Each was transfected separately into a population of U251 and GBM43 PDX cells, with and without LNX1 overexpression. The cells were then cultured for 4 days in order to allow activation of the reporter and were subsequently analyzed by flow cytometry. Results showed significant elevations of Notch1 activity in the LNX1 overexpression population across both vectors and across both cell lines, as measured by GFP or RFP expression in the population (**Fig 5E**). This confirms our hypothesis that LNX1 overexpression results in increased Notch1 activity, which subsequently results in an increasingly stem-like phenotype of the cells.

Finally, after validating that LNX1 overexpression resulted in Notch1 increase, we examined whether LNX1 overexpression promotes stemness in GBM by performing a functional limiting dilution assay in U251 cells as well as the PDX lines GBM43 and GBM6. Our results showed that overexpression of LNX1 significantly increased the stem cell frequency of the population across all three cell lines as stem cell frequency almost doubles in the overexpression condition (*p < 0*.*05;* **Fig 5F**). This result suggests that there is a significant increase in the stemness of the population after overexpression of LNX1.

### Knockdown of LNX1 Activity Results in Reduced Stemness, and Increased Animal Survival

Overexpression of LNX1 shows an expansion of the stem cell compartment. The expansion of the GSC compartment has been closely linked to tumor aggressiveness and poor prognosis for patients [9, 41]. Therefore, we wanted to assess the effect of knocking down LNX1 expression in gliomagenesis. These knockdowns were established using shRNA constructs in the U251, GBM43, and GBM6 cell lines. Lentiviral particles were generated from the plasmids carrying shRNA and cells infected with particles to produce an LNX1 knockdown.

Our first step was to validate our knockdown by western blot. Our results showed that the induction of LNX1 expression in response to TMZ leads to a reduction of Numb. expression and induction of NICD (left panel, scramble shRNA, CT, **Fig 6A**). However, when we knocked down the LNX1 with shRNA, GBM cells failed to reduce NUMB expression in response to TMZ. In response, the level of NICD is significantly lower during TMZ therapy (right panel, LNX1 shRNA, sh4, **Fig 6A**). Furthermore, mRNA levels of Notch1 downstream targets genes RBPSUH, Hes1, and p21expression are elevated as normal following TMZ therapy but are no longer elevated following knockdown of LNX1 (**Fig 6B**). Cells with the knockdown are also shown to be increasingly susceptible to TMZ therapy as compared to control populations (**Fig 6C**).

**Figure 6:**
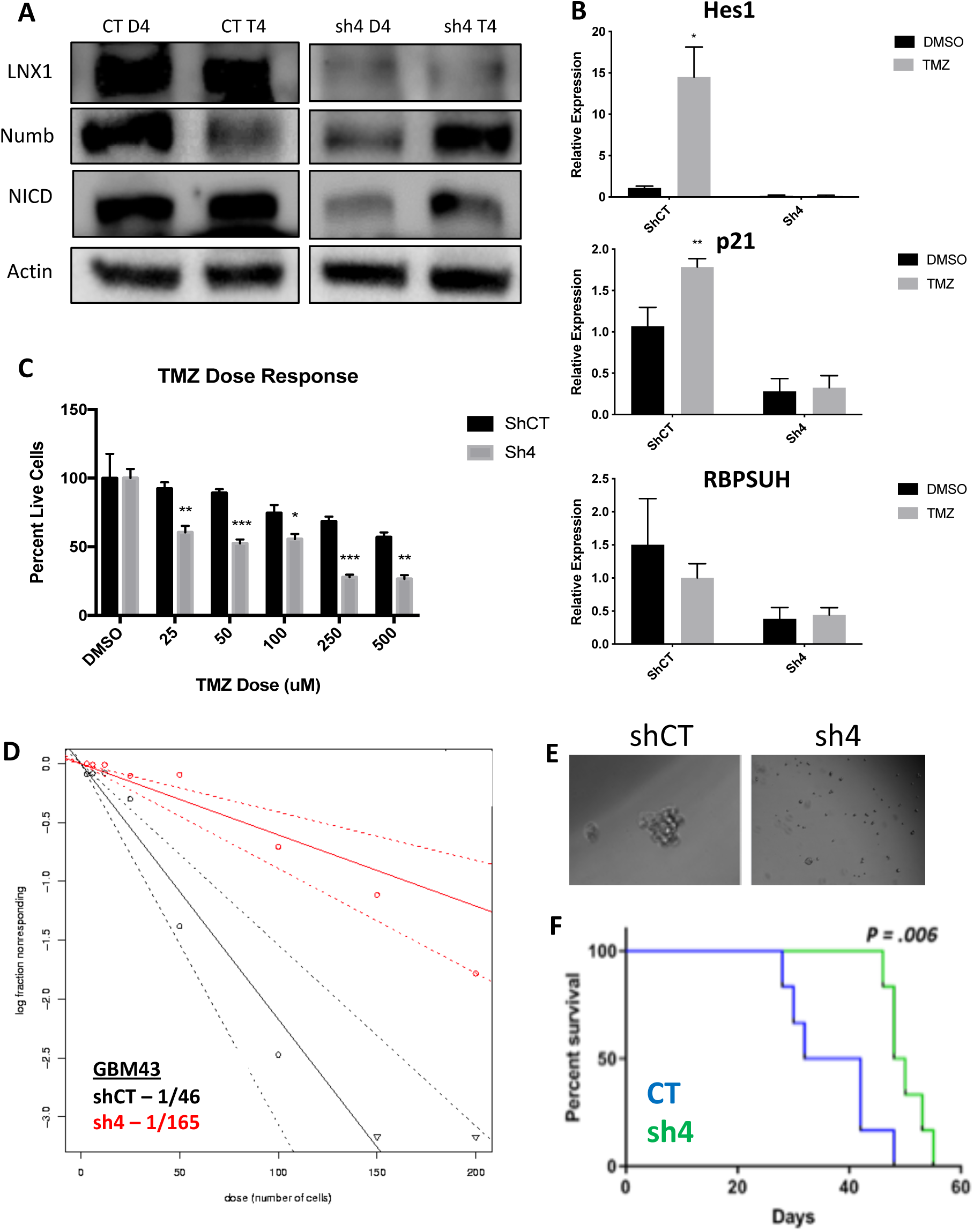
Knockdown Results in Notch Downregulation, Reduced Stemness, and Longer Survival. (A) LNX knockdown results in increased Numb and reduced Notch1 activation. (B) Knockdown of LNX1 resulted in reduced activation of Notch1 downstream markers, as measured by quantitative PCR. (C) LNX1 knockouts were increasingly sensitive to TMZ as compared to controls exposed to the same doses. (D) Limiting dilution assays in showed that the knockdown was no longer able to induce stemness. (E) Representative images showed sphere formation controls but limited sphere formation in our knockdowns. (F) LNX1 knockdown in our animal tumor model showed significant increases in survival over our control population. Statistical analysis was performed in Prism 8, using the Student’s t-test to compare 2 groups or ANOVA for more than 2 groups. Chi-squared testing was used to calculate significance for limiting dilution assays and log-rank testing was used to compare survival curves. *p < 0.05; **p < 0.01; ***p < 0.001; ns, not significant

Next, we examined stemness by a functional limiting dilution assay following LNX1 knockdown by performing a limiting dilution neurosphere assay. In the LNX1 knockdown condition, we noted a reduction in stemness. Visual examination showed a similar trend, with a reduction in neurosphere frequency in the knockdown condition (**Fig 6D**). The stem cell frequency observed in control cells is 1/46, with a decrease to 1/165 after knockdown (*p < 0*.*0001)*. Overall, this supports our hypothesis that knocking down LNX1 results in a significant decrease in stemness, which can result in a less aggressive tumor.

Finally, we performed an *in vivo* experiment comparing our LNX1 knockdown to our control population in the GBM6 PDX line. For this experiment, our previously described animal model of GBM was used. Control or knockdown cells were implanted orthotopically in mice, which are subsequently monitored for the development of clinical symptoms. Mice were sacrificed when it was determined that they were unlikely to survive to the next day by two independent investigators, and their survival times were recorded. Analysis of this experiment showed that the LNX1 knockdown resulted in significantly increased survival times, likely due to the reduced aggressiveness of the GBM cells (*p = 0*.*006;* **Fig 6F**).

## DISCUSSION

GBM is a devastating disease that currently carries a dismal prognosis for patients. As such, it is desperately in need of novel therapies that may provide improved outcomes for patients. Previously, our lab and others have shown that there is a subpopulation of cells in GBM termed GSCs that likely drive the aggressive nature of GBM by driving its therapeutic resistance and recurrence. We have further shown that treatment of tumors with TMZ results in expansion of the GSC compartment, which we believe is occurring due to plasticity-dependent mechanisms within the tumor.

Here, we have shown that cell cycle and stemness programs are significantly upregulated in GBM cells following TMZ therapy. This result has been corroborated by other studies across many different cancers, further bolstering the idea that cell division and expansion of the stem cell compartment are key for tumor progression after therapy [13, 32, 35]. We have additionally shown that polarized cell division regulators are specifically altered after TMZ therapy in GBM, suggesting that the expansion of the stem cell compartment observed after TMZ therapy in part driven by changes in regulators of polarized cell division. Of these regulators, we were able to identify LNX1 as a novel target that is significantly upregulated after TMZ therapy.

Other studies have shown that LNX1 regulates Numb and Notch1 and may have a role in cell division programs [34, 42, 43]. However, no study has examined the specific role of this entire axis and how it may contribute to the expansion of the stem cell compartment, specifically in GBM post-therapy. We were able to show that LNX1 does negatively regulate Numb, which in turn negatively regulates Notch1 expression in GBM cells. In response to TMZ, LNX1 was elevated, resulting in elevations in Notch1 and corresponding gene-level activation of Notch1 downstream genes, which further results in an increased stemness phenotype in cells. This axis was validated both by overexpression studies and by knockdown studies. Furthermore, from a clinical perspective, we were able to show both that LNX1 enrichments do occur in patients and that they are associated with decreases in survival. We were also able to show that loss of LNX1 expression can result in longer survival times in our clinically relevant animal model.

Given that we know GSCs are a major driver of GBM’s aggressiveness, this increased stemness phenotype associated with LNX1 activation suggests that LNX1 may be a promising therapeutic target to block Notch signaling in GBM. Targeting LNX1 may allow for more nuanced modulation of Notch1, which may result in significantly improved outcomes for patients. Notch1 is a well-known modulator of GBM and many clinical trials have been attempted to target Notch1. This is a gene known to be significantly elevated across all GBM tumors and known to be involved in promoting cellular plasticity as it has a multitude of functions in determining cell fate, differentiation, and proliferation. As such, it is able to promote more aggressive cell states. Unfortunately, all therapies developed to target Notch1 have failed due to the unbearable toxicities associated with these inhibitors and the involved dosing schedules required of patients. LNX1 therefore could be a way to more simply target Notch1. Furthermore, since LNX1 is only elevated in GBM cells, it also provides a more nuanced and targeted approach to reducing Notch1 activity.

## CONCLUSIONS

Overall, we believe that this study provides a novel target for treating recurrent GBM. Much of the aggressive nature of GBM has been tied to expansion of the GSC compartment. Preventing this expansion with novel therapeutic strategies could very much change the course of this otherwise devastating disease.

## ACKNOWLEDGEMENTS

This work was supported by the National Institute of Neurological Disorders and Stroke grant 1R01NS096376, 1R01NS112856 the American Cancer Society grant RSG-16-034-01-DDC (to A.U.A.) and P50CA221747 SPORE for Translational Approaches to Brain Cancer. Additionally, this was supported by an internal fellowship (the Medical Student Scholar Program) awarded by the Department of Neurosurgery at Northwestern. The results published here are in part based upon data generated by the TCGA Research Network: https://www.cancer.gov/tcga, and were further analyzed through GlioVis, cBioPortal, and IvyGap.

## AUTHOR CONTRIBUTIONS

<u>Conceptualization</u>: Shivani Baisiwala, Robert H Hall, Atique U Ahmed

<u>Methodology</u>: Shivani Baisiwala, Robert H Hall, Atique U Ahmed

<u>Validation, Formal Analysis, Investigation:</u> Shivani Baisiwala, Robert H Hall, Miranda R Saathoff, Jack M Shireman, Cheol Park, Louisa Warnke, Clare Hardiman, Jenny Y Wang, Chirag Goel, Shreya Budhiraja, Kathleen McCortney

<u>Resources</u>: Atique U Ahmed, Craig M Horbinski

<u>Data Curation & Draft Preparation</u>: Shivani Baisiwala

<u>Review & Editing</u>: Shivani Baisiwala, Atique U Ahmed, Miranda R Saathoff, Jack M Shireman

<u>Supervision, Project Administration, Funding Acquisition</u>: Atique U Ahmed

## SUPPLEMENTARY MATERIALS

**Figure S1:**
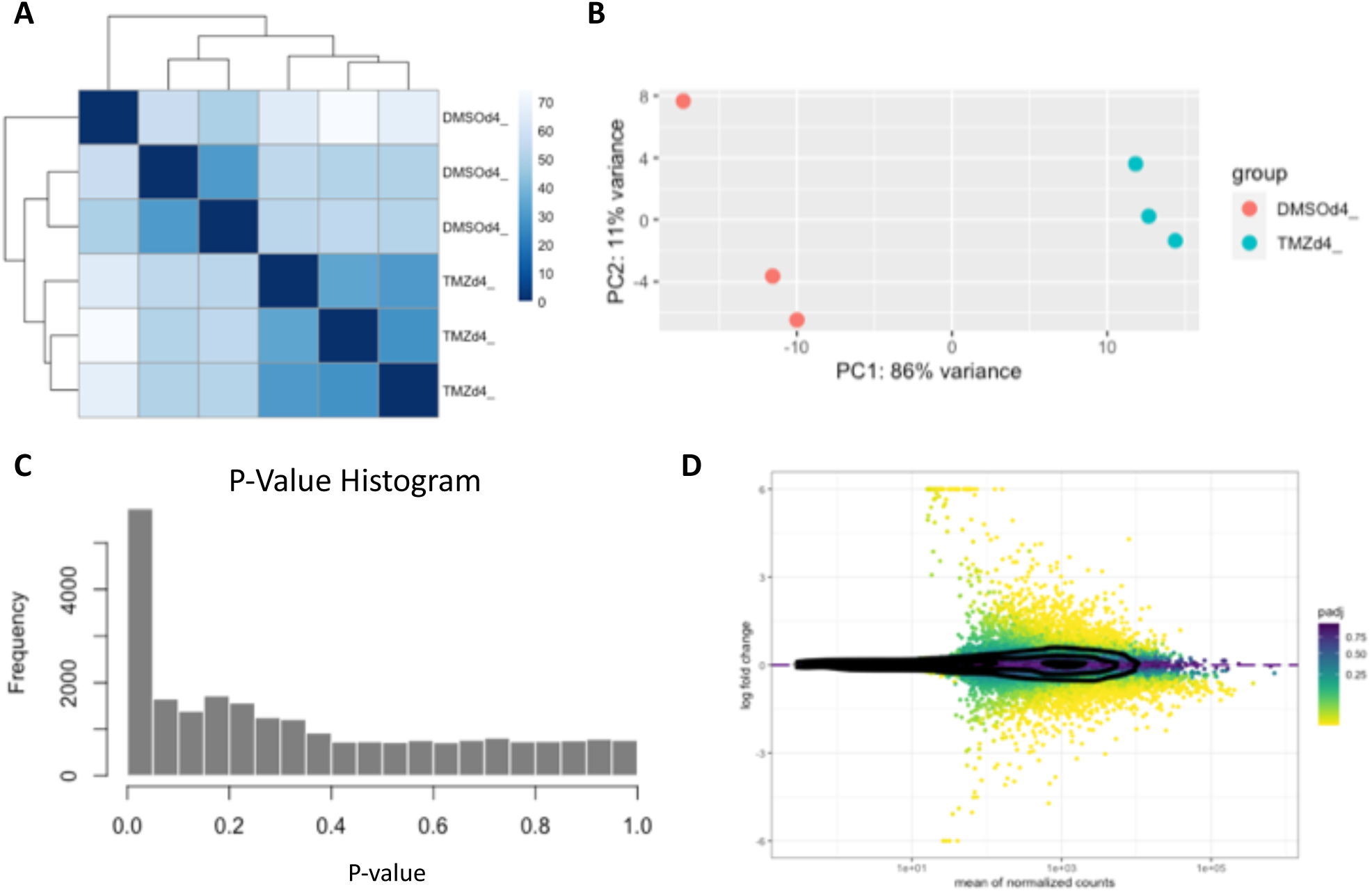
RNA-Seq Quality Metrics. (A) Sample distance clustering shows that replicates are more similar to each other and that there are differences between our control and treatment groups. (B) PCA shows that the greatest element of difference between our samples comes from the treatment performed. (C) Waterfall plot of p-values shows that we have a number of genes that were significantly altered (D) MA-plot represents appropriate shape for trustworthy RNA-Seq data

**Figure S2:**
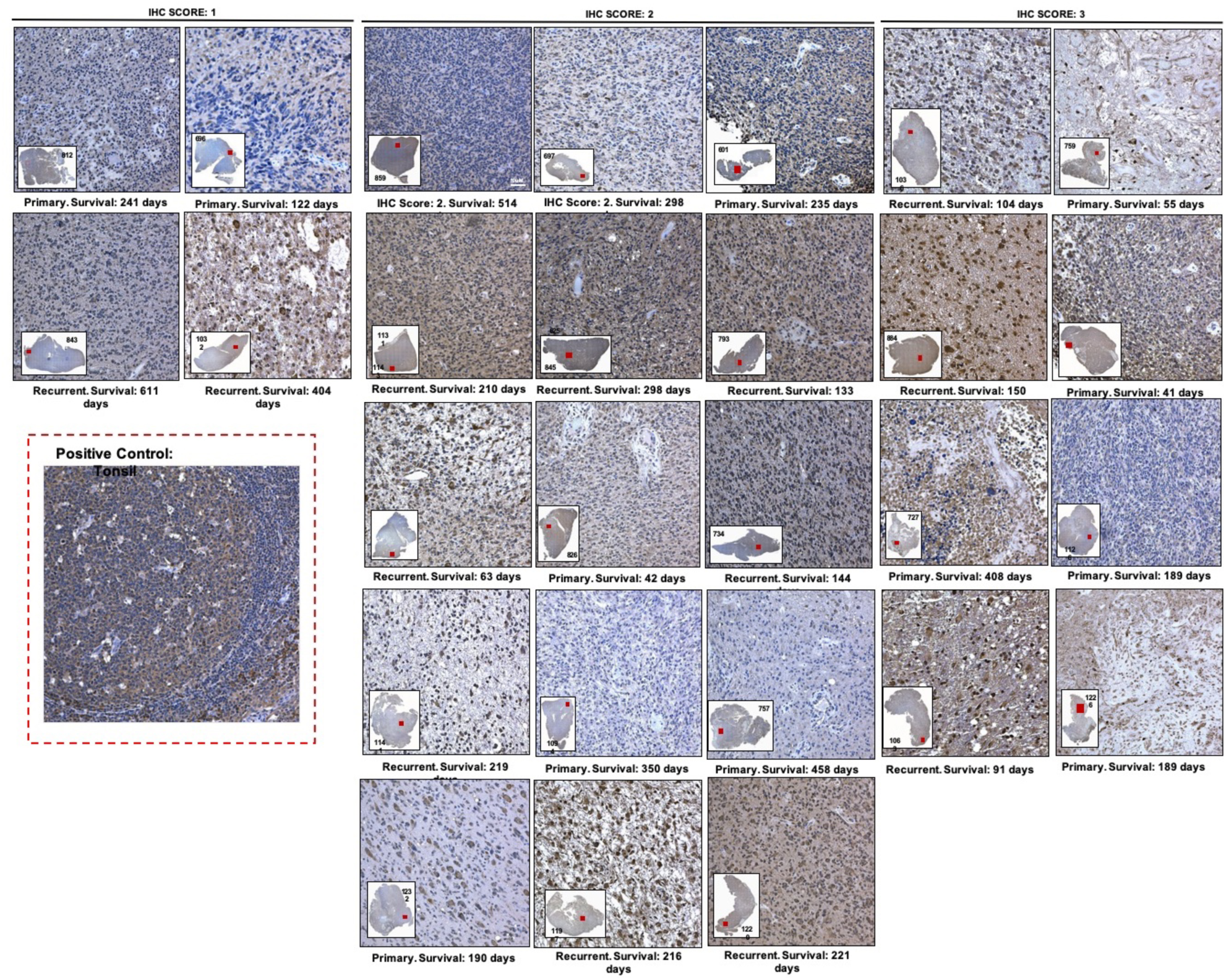
LNX1 Expression in Patient Samples. Images represent immunohistochemistry against LNX1 in GBM patient samples isolated from consenting patients at our institution. Samples were scored by a board-certified neuropathologist and de-identified survival data was correlated with the scores. Positive control for staining was tonsillar tissue, which is known to express LNX1 at high levels.

**Figure S3:**
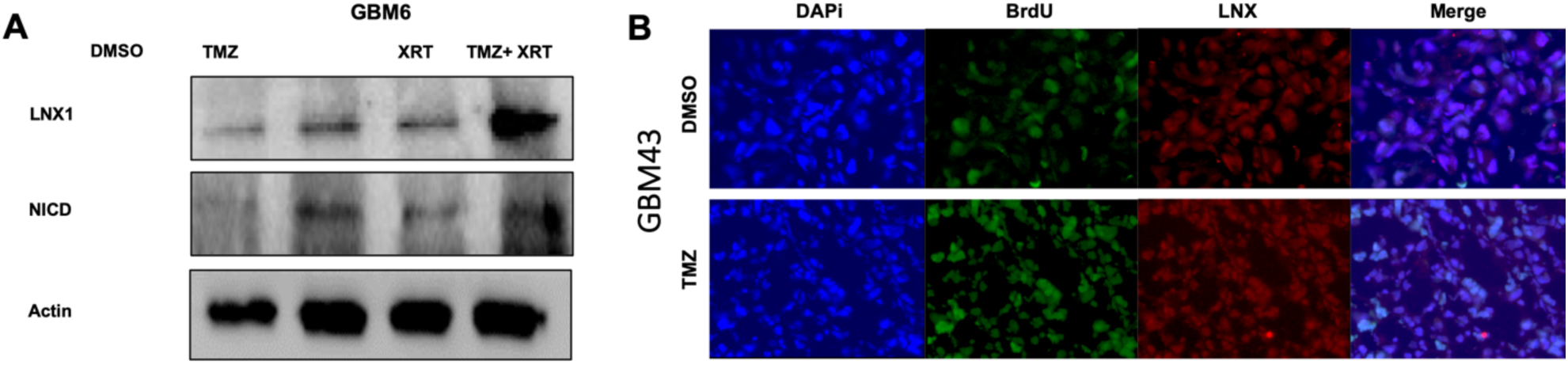
LNX1 Expression Elevates After Exposure to Therapeutic Stress. (A) LNX1 and NICD levels were assessed by western blot after exposure to TMZ, radiation, and combination therapy. Results showed that both were elevated upon exposure to therapeutic stress. (B) GBM43 cells were implanted intracranially to generate an animal tumor model. Tissues were then sectioned and stained for LNX1 and the proliferation marker BrdU. Increased expression of both is noted in tumor areas after TMZ therapy as compared to control DMSO.

**Figure S4:**
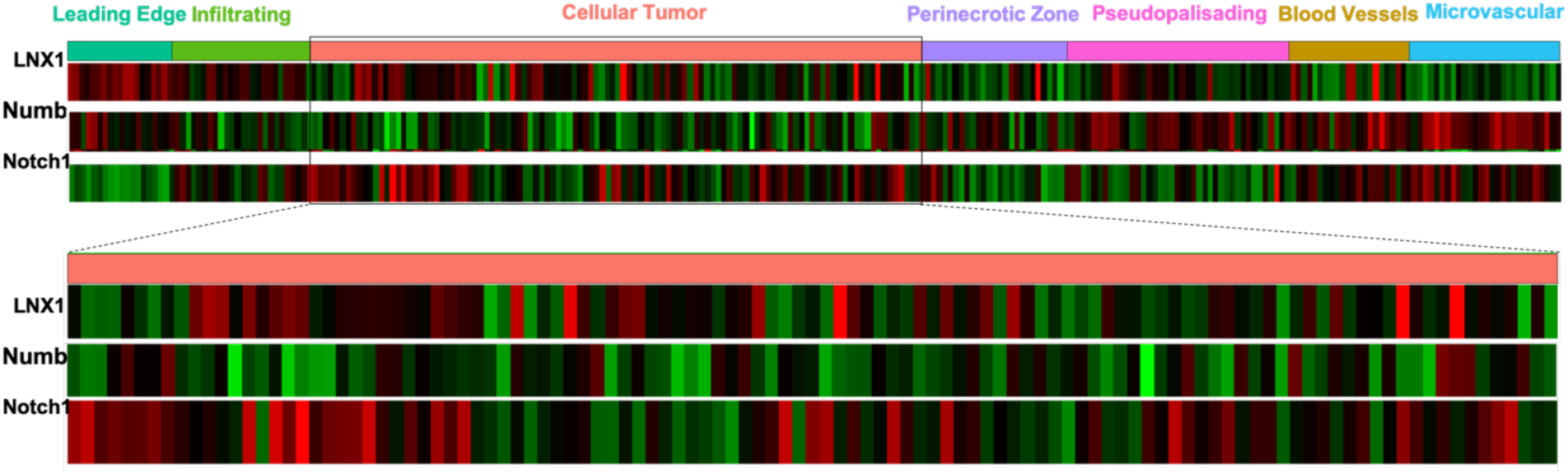
IvyGap provides in situ RNA hybridization data showing expression of genes within various regions of patient tumors. The cellular tumor was assessed for LNX1, Numb, and Notch1 expression and reflected that LNX1 was elevated, corresponding with Numb loss, and Notch1 elevations.

